# metaGE: Investigating Genotype × Environment interactions through meta-analysis

**DOI:** 10.1101/2023.03.01.530237

**Authors:** Annaïg De Walsche, Alexis Vergne, Renaud Rincent, Fabrice Roux, Stephane Nicolas, Claude Welcker, Sofiane Mezmouk, Alain Charcosset, Tristan Mary-Huard

## Abstract

Dissecting the genetic components of Genotype-by-Environment interactions is of key importance in the context of increasing instability and plant competition due to climate change and phytosanitary treatment limitations. It is widely addressed in plants using Multi-Environment Trials (MET), in which statistical modelling for genome-wide association studies (GWAS) is promising but significantly more complex than for single-environment studies. In this context, we introduce metaGE, a flexible and computationally efficient meta-analysis approach for the joint analysis of any MET GWAS experiment. To cope with the specific requirements of the MET context, metaGE accounts for both the heterogeneity of QTL effects across environments and the correlation between GWAS summary statistics acquired on the same or related set(s) of genotypes. Compared to previous GWAS in 3 plant species and a multi-parent population, metaGE identified known and new QTLs. It provided valuable insight into the genetic architecture of several complex traits and the variation of QTL effects conditional to environmental conditions.

## 1 Introduction

Understanding the adaptation mechanisms of plant species to different environments and the underlying genetic architecture has been a long-standing challenge in plant genetics [20, 4, 15]. It can be investigated through the mapping of quantitative trait loci (QTL) and the evaluation of their effects in different environments. Multi-Environment Trials (MET) that consist in phenotyping the same panel of genotypes in different locations and/or over different years have been widely adopted to dissect the genetic components underlying the Genotype-by-Environment (G×E) interactions and to describe the QTL response to different environmental factors. Recent genome-wide association studies in plant genetics have illustrated how the QTL-environment interaction may affect the phenotypic response [41, 50, 1, 18]. While informative and promising, QTL identification through the analysis of MET requires a dedicated statistical methodology to account for both the instability of allelic effects and the high level of correlation between measurements acquired on the same (or very similar) panel(s) across environments. Compared to the single environment analysis scenario for which many computationally efficient and grounded methods have been developed [57, 59, 36, 37, 32], the statistical analysis of MET is still an open problem with only a few methods available [31, 60, 7, 38]. Powerful and scalable methods that account for the environmental variation and its interaction with the QTLs remain challenging.

In this context, an appealing alternative to classical approaches consists in performing individual GWAS analyses in each environment separately using one of the efficient single environment methods mentioned above, then jointly analyzing the summary results - typically the effects and p-values associated with the different markers - of the individual GWAS through a Meta-Analysis (MA) [17, 19]. GWAS-MA has proven to yield significant gains of power over initial individual analyses while efficiently controlling for false positives and keeping the computational burden low [13]. It has been successfully applied to both human [8, 44, 34] and animal [10, 25] genetic studies, allowing the detection of QTLs with small or moderate effects[28]. GWAS-MA was, however, rarely applied to plant genetics [58, 51] and never in the context of MET analyses. In addition, most MA procedures have been developed for human genetics purposes and aim at detecting QTLs with stable effects over independent populations and, as such, are not suited to the MET context where important variations in QTL effects can be observed and where the same (or related) panel(s) is(are) phenotyped across different locations and years.

We propose new extensions to make MA procedures amenable to MET and G×E interaction analysis in plant genetics. We developed fixed and random MA procedures to handle respectively fully controlled environments, and experiments where the monitoring is insufficient to describe and classify the environments, as often observed in fluctuating field conditions. To further investigate the G×E interaction, we introduce a new test procedure to identify QTLs whose effect variations are correlated with a given environmental covariate. We demonstrate the efficiency of our approach through applications to MET GWAS analysis of 3 species (A *thaliana*, maize and wheat) and a maize multi-parent population MET. The whole statistical procedure is available in the metaGE R package.

## 2 Results

### 2.1 Meta-analysis approach

The study of G×E interactions in plant genetics requires the evaluation of the same panel - or highly overlapping panels including several common genotypes - evaluated under different environmental conditions. Depending on the experiment, environments may correspond to controlled stress conditions (*e.g*. nitrogen, water or competition stress), or to different fields and/or years where the environmental conditions are contrasted but not fully controlled by the experimenter. The newly developed fixed effect (FE) and random effect (RE) meta-analysis procedures respectively cope with the controlled and uncontrolled environment cases. They are briefly described (see the section 4.2 for details), then illustrated on several applications cases.

We consider a meta-analysis relying on *K* different GWAS performed in individual environments testing the association between a set of *M* markers and a phenotype of interest. We denote *β_mk_* the estimated effect of marker *m* in environment *k*, and *p_mk_* is the associated p-value. We define the z-score *Z_mk_* as

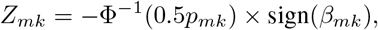

where Φ stands for the standard Gaussian cumulative distribution function. The smaller the p-value of the marker, the greater will be the absolute value of the z-score. The sign of the z-score corresponds to the sign of the marker effect. Importantly, when a marker *m* is not associated with the phenotype, *Z_mk_* follows a standard Gaussian distribution.

#### Fixed Effect (FE) procedure

When the environmental conditions are controlled, the environments can be *a priori* classified into several groups. One can then assume the marker effect to be stable within each group but different from one group to another. Assuming the environments are classified into *J* distinct groups, the fixed effect model for a given marker *m* is:

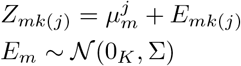

where 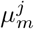 is the marker effect specific to group *j* = 1, *…, J*, *E _m_* = (*E_m_*_1_*, …, E_mK_*) is the vector containing the residuals error, and Σ is the inter-environment correlation matrix. Testing the association at marker *m* corresponds to testing the null hypothesis of all group-specific marker effects 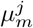, *j* = 1, …, *J* being equal to zero.

#### Random Effect (RE) procedure

In the case of uncontrolled or partially controlled environmental conditions, the heterogeneity of the QTL effects across environments may be accounted for through a random marker effect. The model is updated as follows:

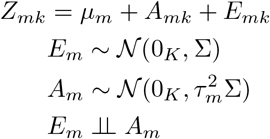

where *μ_m_* is the mean marker effect, *A_m_* = (*A_m_*_1_*,…, A_mK_*) is the random effect accounting for the heterogeneity of the marker effect, with variance 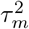. Testing the association of the marker *m* corresponds to testing the null hypothesis of the mean marker effect *μ_m_* and the marker effect variance 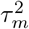 being equal to zero.

To further investigate the G×E interaction, we also introduce a new meta-regression test procedure to identify QTLs whose effects variations are correlated to a given environmental covariate (described in Section 4.2).

### 2.2 Applications

The 3 datasets presented in the following sections were selected to represent classical MET experiments for *G × E* interactions association studies in plant quantitative genetics, and cover controlled or partially/not controlled experimental designs and a multi-parent MET population. Additionally, a fourth application to a wheat MET GWAS dataset is presented in Supp. Mat. A. All datasets are publicly available (see the Data Availability section 4.3).

#### Controlled experiments

We consider the Arabidopsis dataset of [23], where GWAS analyses were performed on the local mapping population TOU-A, with a panel of 195 whole-genome sequenced accessions evaluated in six controlled micro-habitats (combinations of three soils × presence/absence of inter-specific competition, noted A to F). Each accession was phenotyped for bolting time and had genotypic information for 981,278 SNPs (after quality control and a MAF threshold of 0.07). The trials were conducted under controlled stress conditions, with Arabidopsis being grown in competition with the weed *Poa annua* in environments B, D, F and without competition in environments A, C and E. The fixed-effect procedure described in details in the Methods Section 4.2 was applied to perform the joint analysis of the 6 GWAS summary statistics.

First, a standard FE procedure was performed to detect markers with a stable effect across environments, leading to the identification of 191 SNPs clustered into 61 QTLs. Among these QTLs, 51 were found significant in at least one individual GWAS (see Table S1 in Supp. Mat. B for details). Following [11], the list of significant SNPs was further validated by looking at the enrichment ratio for *a priori* candidate genes in bolting time. The enrichment of the identified markers (enrichment=4.13) was significantly higher than what would be expected at random ([*q*_0.05_; *q*_0.95_] = [0.066; 3.2]).

We then applied the contrasted FE testing procedure described in the Methods section, Equ. (2) with a group effect corresponding to the presence or absence of a competition with *Poa annua* to detect markers with contrasted allelic effects based on the presence or absence of competition. The contrasted FE procedure identified 221 SNPs located in 72 QTLs (Fig.1.A) and covering 160 candidate genes that show significant enrichment for three Gene Ontology terms (MapMan functional annotation,[46]), *i.e*. ‘development’ (P = 8.866e-03), ‘cell’ (P = 1.522e-03) and ‘tetrapyrrole synthesis’ (P = 0.020). Interestingly, the two latter MapMan processes were also detected as enriched when challenging the local mapping population TOU-A to the presence of three weed species in greenhouse conditions [35].

**Figure 1:**
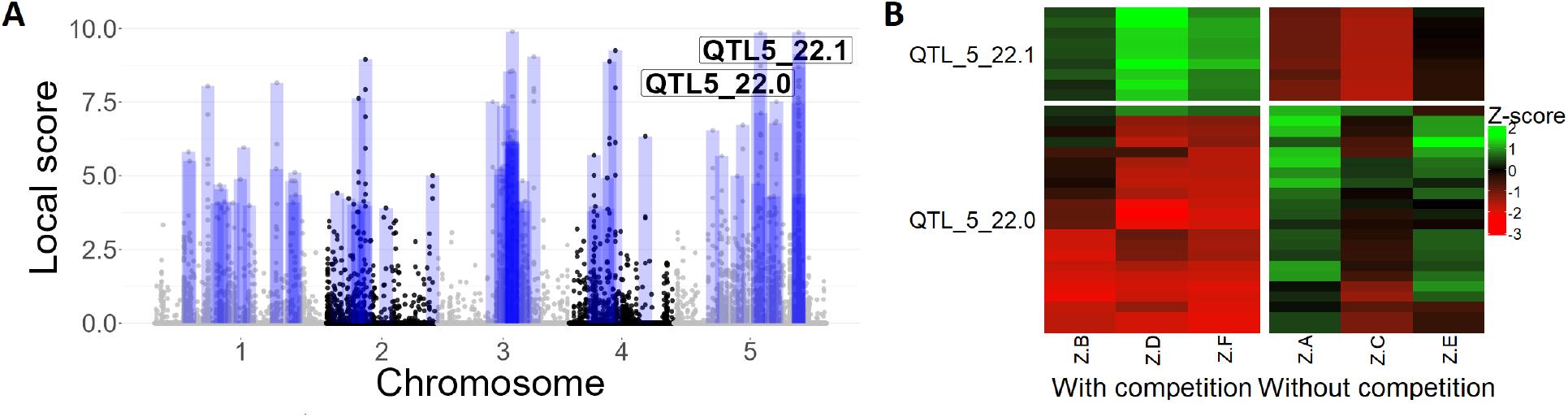
Results of the FE procedure applied to the Arabidopsis dataset to detect markers with competition contrasted effects. (A) Local scores along the chromosomes. The boxes represent the significant zones identified. (B) Z-scores of two QTL regions located on chromosome 5 (QTL5_22.0 a,d QTL5_22.1), with markers in rows and environments in columns. A, C and E correspond to the 3 environments without competition.

Fig.1.B represents the z-scores of two QTLs found on chromosome 5 involving 22 and 9 markers, respectively. The identified markers clearly exhibited a contrasted marker effect profile: when the effects were positive in one of the two groups of environments, they were negative or null in the other. All the 22 SNPs of the second QTL (QTL5_22.1) are located in the AtCNGC4 genomic region that is well-known to affect floral transition [12] [22].

Importantly, as the contrasted FE procedure aims at identifying markers with contrasted (*i.e*. unstable) effects across the two sets of environments, 71 out of the 72 QTLs identified by the contrasted FE procedure are new candidates that were not detected by the standard FE procedure.

#### Uncontrolled experiments

We consider the Maize dataset of [41], where GWAS analyses were performed on a panel of 244 maize dent lines evaluated as hybrids with a common parental line (a usual practice in maize genetics) in 22 environments (combinations of location × year × treatment). Each line was genotyped at 602,356 SNPs (after quality control) and phenotyped for grain yield (GY) in the 22 environments. In addition, environmental variables were measured in each environment.

The RE procedure detailed in Section 4.2 was applied to perform the joint analysis of the 22 per environment GWAS summary statistics. In total 52 genomic regions were identified, of which 14 correspond to QTLs also detected in the original publication [41]. The three QTLs with the most significant association peaks (Fig.2.A) were located on chromosomes 3 (QTL3_120.0) (local score=38), 6 (QTL6_20.3) (local score=415) and 7 (QTL7_41.4) (local score = 18, not detected in the original publication). The z-score heatmaps corresponding to QTL6_20.3 and QTL7_41.4 are displayed in Fig.2.B. The heatmap of QTL6_20.3 revealed a cluster of six environments (Cra12R, Cam12R, Cra12W, Mur13R, Mur13W and Cam12W) with strong similar marker effects that were found to be characterized by severe heatwaves with high night temperature (close to 22°C) and high maximal temperature (above 36°C) together with high evaporative demand during the day (3.6 KPa). The QTL6_20.3 individual GWAS p-values were all lower than 1e-6 in the 6 aforementioned environments, confirming the strong association signal detected by the RE procedure. In contrast, the QTL7_41.4 showed a moderate positive effect across nearly half of the environments: individual GWAS p-values were lower than 1e-2 in ten environments out of 22, and was found significant in only two individual GWAS.

**Figure 2:**
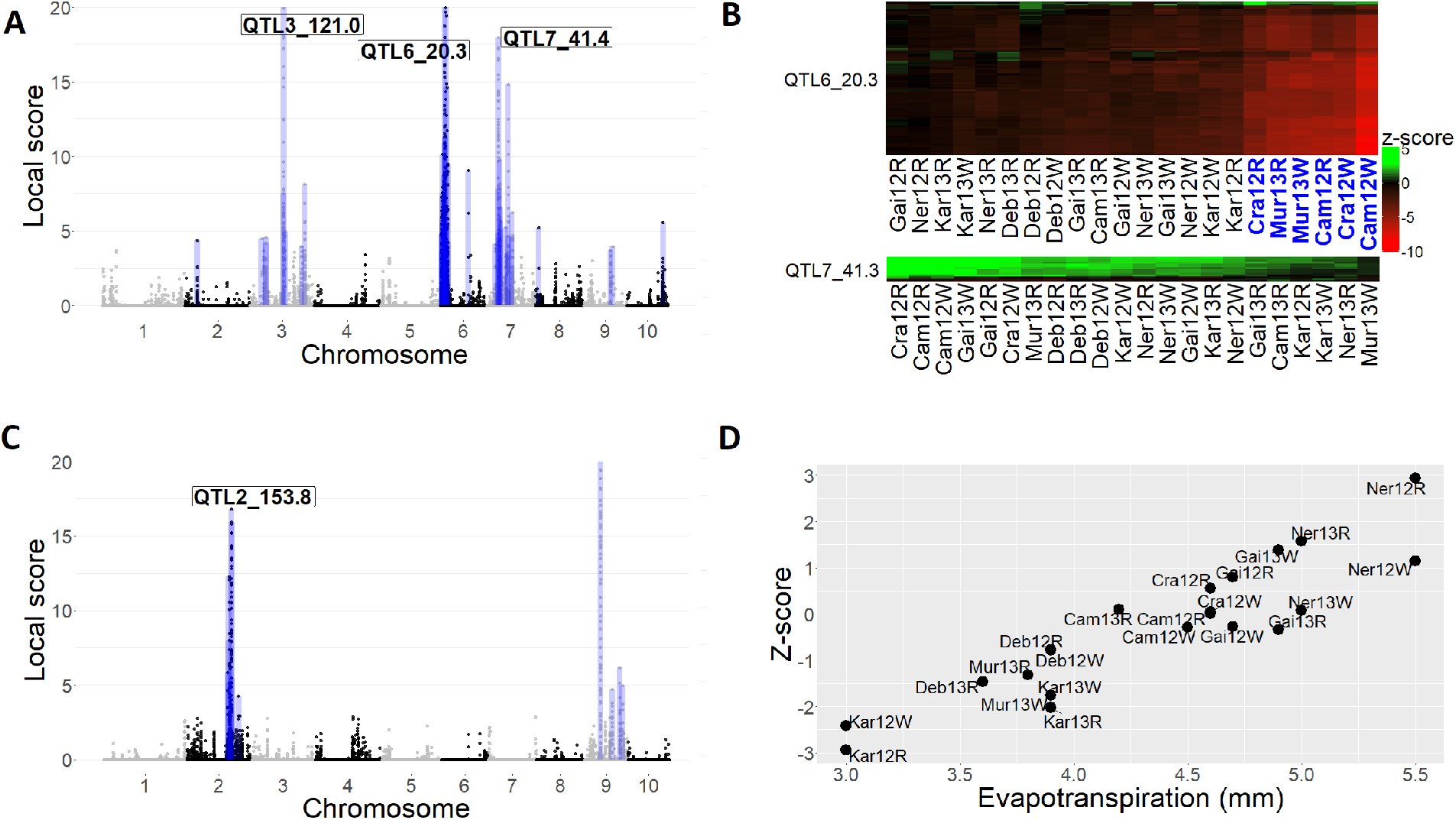
Results of the MA procedure applied to the Maize dataset. (A) Local score obtained from the RE procedure along the chromosomes. The range of values on the y-axis has been bounded to 0–20 to highlight minor QTL. The boxes represent the significant zones identified. (B) Z-scores of two QTLs located on chromosomes 6 and 7 (QTL6_20.3 and QTL7_41.4), with markers in rows and environments in columns. Only the 100 top significant SNPs out of 169 composing the QTL6_20.3 have been displayed. The six environments highlighted in blue correspond to environments characterized by severe heatwaves (night temperature close to 22°C and maximal temperature above 36°C). (C) Local score obtained from the meta-regression test for the evapotranspiration along the chromosomes. The range of values on the y-axis has been bounded to 0–20 to highlight minor QTL. The boxes represent the significant zones identified. (D) Z-scores as a function of the evapotranspiration (ET0), for the top significant marker of QTL2_162.5 detected with the meta-regression procedure (Maize dataset, marker AX-91538480).

We performed a genome-wide detection of markers whose effect variations are correlated with environmental variables by running the meta-regression test procedure described in Section 4.2. The environmental variables considered were the evapotranspiration (ET0), the mean night temperature during the flowering period (Tnight) and the mean night temperature during the grain filling period (Tnight.Fill).

Regarding evapotranspiration, the meta-regression test identified 14 QTLs located on chromosomes 2 and 9 (Fig.2.C). Allelic effects on grain yield of one of the most significant marker (marker AX-91538480, QTL2_153.8) markedly changed with potential evapotranspiration (ET0) (Fig.2.D).

For the mean night temperature during the flowering period, the meta-regression test identified 21 QTLs, with the main association corresponding to a genomic region located at less than 0.6 Mb to QTL6_20.3 (Fig.S2.A in Supp). This detection corroborates the results previously obtained in [41]).

Regarding the night temperature during the grain filling period, the meta-regression test identified 15 QTLs across six chromosomes. In particular, the allelic effects on GY of one of the most significant marker located on chromosome 9 (marker AX-91123283, QTL9_28.6) changed dramatically according to night temperature during the grain filling period, with positive effects on cool night and negative effects on hot night during the grain filling period (Fig.S2.B in Supp). See Supp.Mat C for an analysis regarding night temperature during the grain filling period.

#### Extension to Multi-Parental Population MET experiments

The maize EU-NAM Flint includes 11 biparental populations obtained from crosses between UH007 and 11 peripheral parents representative of the Northern Europe maize diversity [5, 33]. In each population, double haploid (DH) lines were produced and genotyped at 5,263 SNPs (after quality control). All populations were evaluated for biomass dry matter yield (DMY) in 4 locations (La Coruna, Roggenstein, Einbeck and Ploudaniel). Three populations with less than 30 progenies were removed from the present analysis. Unlike previous datasets involving an association panel, the present one consists in a multi-parent crossing design, each bi-parental progeny being phenotyped in all aforementioned locations. In this context we extend the notion of environment to the combination of one sub-population and one location.

To estimate the allelic effect per parent and environment, GWAS analyses were performed on each combination of cross and location, resulting in 32 individual analysis. The RE procedure detailed in Section 4.2 was applied to perform the joint analysis of the 32 individual GWAS summary statistics. In total, 16 QTLs were identified, highlighting some very significant association peaks, especially on chromosomes 1 (QTL1_117.6) and 6 (QTL6_84.2). These QTLs were also identified in the publication of Garin et al. [24]. The allelic effect of QTL1_117.6 was almost consistent across all populations except F2 (Fig.S3 in Supp).

QTL6_84.2 showed an interesting genetic effect series since an ancestral allele inherited by parents D152, F03802, F2, F283, UH006, and DK105 [24] had a strong negative effect mostly in the environment TUM (Fig.3.A).

**Figure 3:**
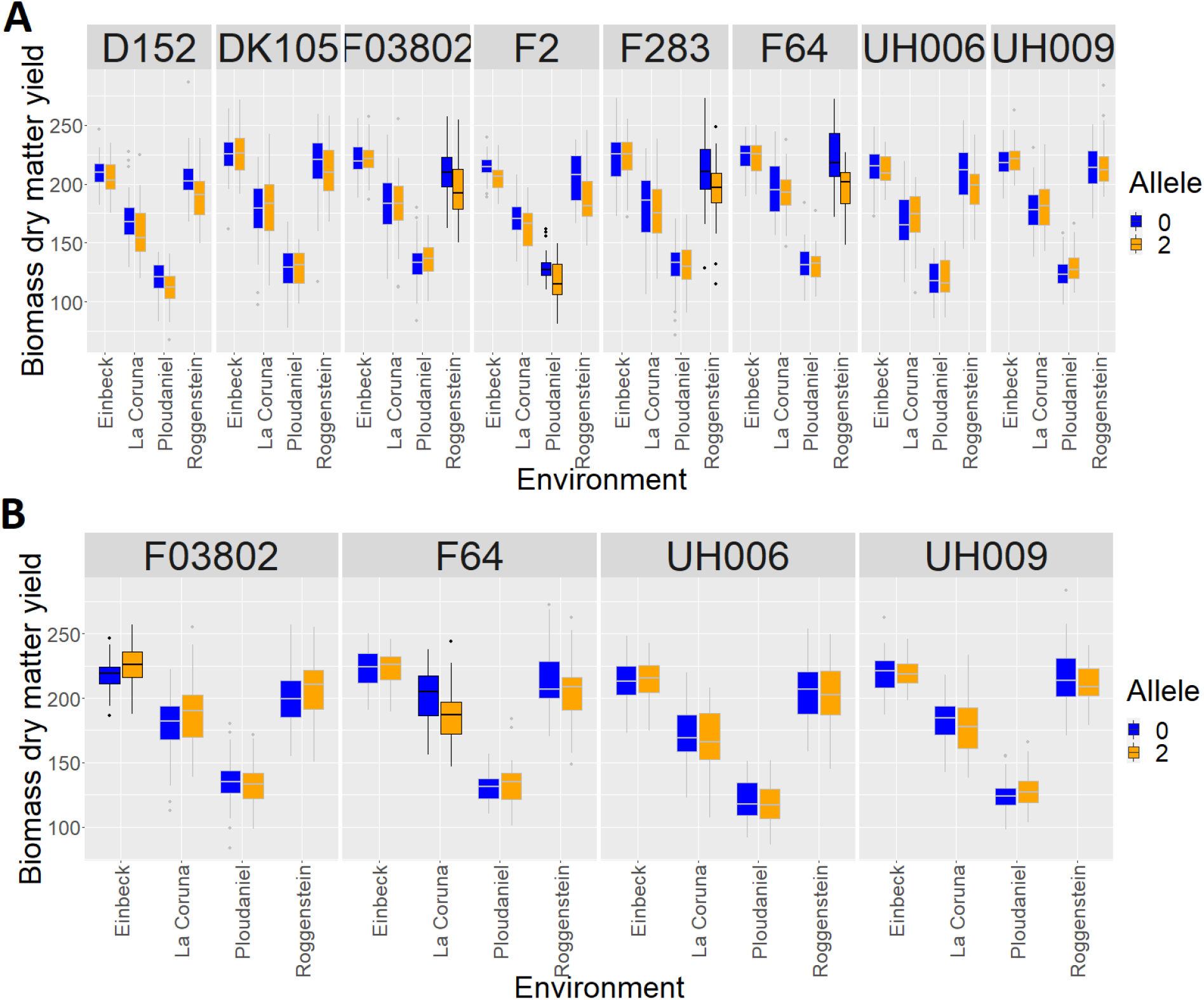
(A) Allelic effects of QTL6_84.2 (EU-NAM Flint, marker PZE.106101278) through locations and sub-populations (B) Allelic effects of QTL5_23.9 (EU-NAM Flint, marker PZE.105012387) through locations and sub-populations. A combination of location x sub-population is highlighted if individual GWAS p-value were below 0.01.

While the analysis of Garin et al. [24] was limited to the study of two of the four locations, the present analysis included all locations. In this study, we revealed ten new QTLs not detected in the initial analysis, five of them correspond to QTL which showed effect inversion among populations. For example, the QTL5_23.9 showed a positive effect for the population F03802 and a negative effect for the population F64 (Fig.3.B). We also noticed that 3 of them corroborated with flowering time QTL detected from the same materials in Giraud et al. [26]. Flowering time has a simpler genetic determinism than grain yield and is one of its main drivers, with negative, null or positive correlations according to environmental conditions [41].

#### Computational efficiency of the method

The computational times corresponding to the meta-analysis procedure for each dataset are displayed in Table 1.

**Table 1:** Computational time of the whole meta-analysis procedure (in detail, the time of the inference of the inter-environment correlation matrix) for each dataset.

| Dataset | Nb. Env. | Nb. Markers | Method | Time |
| --- | --- | --- | --- | --- |
| Arabidopsis | 6 | $\approx 1M$ | FE | 1.2mn (26s) |
| Maize | 22 | $\approx 600K$ | RE + MR | 2.25mn (41s) |
| Wheat | 16 | $\approx 100K$ | RE | 47s (30s) |
| EU-NAM Flint | 32 | $\approx 6,000$ | RE | 12s (8s) |

The analysis of the different datasets considered in this article was handled in less than 3 minutes, even for datasets characterized by a large number of environments and/or markers. Importantly, most of the computational time corresponded to the inference of the inter-environment correlation matrix, a step that needs to be run only once as it is common to both the RE and FE models and all subsequent testing procedures.

## 3 Discussion

We proposed new key extensions to make GWAS meta-analysis (MA) amenable to MET analysis and G×E exploration in plant genetics. This made it possible to address a need in the community that we illustrated through four case studies in A *thaliana*, maize and wheat, which cover a diversity of experimental designs and trait complexity.

GWAS-MA has proven to yield significant gains of power over initial individual analyses while efficiently controlling for false positives [13]. The observed gain in detection power applies as well to the MET context. Our MA approach revealed interesting new QTLs even in regions where the test statistics did not pass the nominal significance threshold in most individual environments. For example, the genomic region corresponding to the third highest local score in the Maize dataset (QTL7_41.4, local score = 18) was found significant in only two environments out of 22 and was not detected in the original publication [41]. This region harbors QTLs controlling plant growth rate and final biomass in water deficit conditions [45]. Similarly, in the Wheat dataset none of the QTLs identified by our MA RE method were found significant in any environment. Therefore, the joint analysis of individual GWAS is suitable for complex traits (such as yield) whose genetic variations are usually due to many QTLs with minor effects that might go undetected in single environment analyses.

In addition to the gain of power, our MA method allows testing any contrast between the effects of environmental subgroups. Although tests of contrasts have been formerly considered in the MA context [9], it was restricted to the comparison of effects across subgroups and to the case of independent individual studies. The extension presented here facilitates the assessment of QTL effects stability or variation across environmental conditions, which is of key interest to detect alleles conferring specific adaptive features. As an illustration,the analysis of the Arabidopsis dataset highlighted a major region on chromosome 5 (QTL5_22.0), with an effect sign that switched according to the presence or absence of competition with *Poa annua*. Arabidopis QTL5.220 is located in the AtCNGC4 genomic region that is well-known to affect floral transition [12, 22]. In addition, AtCNGC4 impairs plant immunity [22, 54], which is in line with the negative effect of competitive interactions on pathogen defense to the benefit of plant development such as floral transition [53].

Similar results were found in the Maize analysis where environmental conditions were not controlled a priori. The analysis of allelic effects of QTL6_20.3 detected with the RE model highlighted a group of six environments (Fig.2.C). A subsequent analysis showed that these environments were characterized by severe heatwaves at night. Consistently, QTL6_20.3 colocalized with a QTL affecting grain yield in the “Po valley” in which maize is currently irrigated but subject to high temperatures during summer [16]. QTL6_20.3 also overlaps with a large 2.4 Mbp Present/Absent Variant (PAV) harboring dozen of genes - including one encoding an ABA-induced protein by water deficit [41]. These genes were further shown to be associated with environmental adaptation to high temperature and to have undergone strong selection during both domestication and improvement [29].

Beyond *a posteriori* interpretation, environmental variables can also be incorporated as a covariate in the GWAS-MA model for a whole genome scan of the response of QTL effects. Such a relationship between marker effects and environmental variables was investigated in [41] for the Maize dataset. However, this initial analysis was performed empirically (*i.e*. without any proper testing procedure) and was restricted to markers found significantly associated in at least one environment. We reanalyzed the full dataset using our Meta regression procedure using 3 covariates and identified a number of new regions. Figure.2.D illustrates how the effect of a QTL located on chromosome 2 (pos154) varies linearly from negative to positive effects according to evapotranspiration. This QTL colocalizes with the QTL of expression of aquaporins (eQTL of PIP2.2 ez and eQTL of PIP2.1 ez). These channel proteins facilitate water transport between cells and impact water use efficiency (WUE) and stomatal conductance (Granato et al., in prep). These two physiological adaptive traits are highly sensitive to environmental conditions [45]. A second region was detected on chromosome 9 (pos125) using the same environmental covariate. It co-localizes with a QTL of plant growth (PG) rate and a QTL of WUE under water deficit conditions. Both PG and WUE are two traits that are highly sensitive to evaporative demand. Lastly, a third QTL, also located on chromosome 9 (pos135), co-localizes with a QTL of plant growth sensitivity to soil water potential, consistent with observed effects [45]. None of these genomic regions on chromosomes 2 and 9 were detected in the initial publication.

An attractive feature of MA procedures is their ability to cope with unbalanced/incomplete data - without need for further data imputation or additional computational overhead. The procedures presented in this article rely on summary statistics (*i.e*. p-values and effect signs) obtained from per-environment GWAS, which avoids re-scaling if traits were measured on different scales. The use of summary statistics also makes the addition/removal of a given environment (based *e.g*. on *post hoc* quality control) and the update of the results straightforward. Furthermore, MA procedures do not require the availability of phenotypic evaluations for all individuals in all environments. Regarding the genotypic information, MA procedures easily handle cases where summary statistics are missing for some markers in a series of environments. This may occur *e.g*. when different technologies or sequencing depth were used in the individual experiments. Furthermore, in plant genetics, Multi-Parental Population analyses are pervasive and also yield missing summary statistics as different sets of markers may be mono-morphic in different parental sub-populations. Our method is still applicable in such cases as illustrated with the EU-NAM example where inversions of allelic effects were identified for some sub-populations (Fig 3). These inversions may correspond to a genetic background effect (*i.e*. to conditional epistasis, see for instance [6]), or alternatively to an allelic series with several haplotypes presenting a range of effects, with the SNP allele contrasting central haplotypes vs. extreme opposite haplotypes.

While several approaches have been proposed to model the G×E interactions in different contexts [31, 60, 7, 38, 24], many of them exhibit a computational time that may become prohibitive when dealing with either large scale genomic datasets or a large number of environments. In contrast, the G×E MA procedures presented here do not suffer from such limitations. As an example the QTLs detection analysis of the EU-NAM Flint (5,000 markers) required 15 seconds using the MA procedure, and more than 7 hours using the MPP-ME methodology of Garin et al. [24]. More generally the different panels considered in this article were handled within minutes, most of the computational time corresponding to the inference of the inter-environment correlation matrix (see Table 1).

Our methods can be applied to various MET designs such as controlled experiments, uncontrolled or partially controlled experiments and all kinds of multi-parent populations MET widely used in modern breeding programs. Our methodology integrates several tools for the characterisation of environments. The contrast test associated to the FE procedure enables the identification of QTLs specific to some subsets of environments, *i.e*., whose behaviour interacts with the environmental characterisation defined by the classification. The definition of the classification is very flexible and can rely on any qualitative covariate, such as known control/stress conditions or sub-populations in the case of an MPP experiment. The meta-regression test allows the detection of QTL whose variations are correlated to any quantitative environmental covariate. Searching for such relationships between marker effects and environmental characteristics is a key issue in plant genetics. It has also been widely investigated in the context of of animal and human MA-studies, where dedicated procedures have been developed to handle environmental covariates measured at the individual level [3, 39]. Our meta-regression procedure represents a new contribution to account for environmental covariates measured at the environment level, and complement the existing MA toolbox. All the testing procedures presented here, along with the FE and RE procedures are implemented in the metaGE R package available on the CRAN repository.

In the recent years, a number of public initiatives involving thousands of individuals evaluated in dozens of well characterized environments have been developed, such as the Genomes To Fields project for maize [2] or the elite yield trial nurseries from CIMMYT’s bread wheat breeding program [30, 43]. MET are one of the central elements of a breeding program [14]. Screening such datasets for genome-wide associations requires the development of scalable and flexible methodological tools. In this context G×E Meta-Analysis can be routinely applied in most breeding programs and we strongly believe that it will represent a methodology of choice in the future to address the many challenges of modern GWAS.

## 4 Methods

### 4.1 Meta-analysis classical approach

In this section, we present the fixed effect and the random effect meta-analysis that are classically used in human genetics [52] [17]. The newly developed methods necessary for its implementation on MET in plant genetics will be presented in Section 4.2.

#### Fixed effect procedure

##### Model

We consider a meta-analysis relying on *K* different genetic association studies testing the association between a set of *M* markers and a phenotype of interest. We denote *β_mk_* the estimated effect of marker *m* in study *k*, and *p_mk_* its associated p-value. We define the z-scores *Z_mk_* as

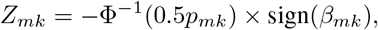

where Φ^*−*1^ stands for the standard Gaussian cumulative distribution function. The z-score is to be understood as follows: the smaller the p-value of the marker, the greater the absolute value of the z-score, with the sign of the z-score corresponding to the sign of the marker effect. Importantly, when marker *m* is not associated to the phenotype, *Z_mk_* follows a standard Gaussian distribution, *i.e*. the *H*_0_ distribution of *Z_mk_* is known. The fixed-effect model assumes the effect of a marker to be stable (*i.e*. identical) across studies. Denoting *Z_m_* ≔ (*Z_m_*_1_*,…, Z_mK_*) the vector of z-scores of marker *m*, one has

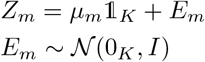

where 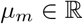 quantifies the deviation to *H*_0_ of marker *m* and is common to all studies, *E_m_* the vector of error terms, and *I* the identity matrix. Note that the model assumes the z-scores to be mutually independent. This assumption is satisfied whenever the initial GWAS analyses are performed on different panels, a classical configuration in human genetics where MA is usually performed to summarize GWAS performed on different populations.

##### Inference

The parameter *μ_m_* can be easily inferred using the empirical mean of *Z_m_*:

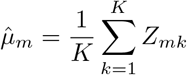

One can then perform association detection by testing

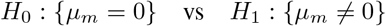

based on the following test statistic

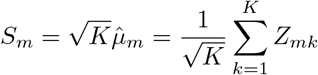

that follows a 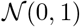 distribution under the null hypothesis, which corresponds to the approach of the METAL procedure [56]. The resulting MA p-value for marker *m* is then

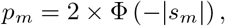

with *s_m_* the observed value of *S_m_*.

#### Random effect procedure

##### Model

We denote *β_mk_* the estimated effect of marker *m* in study *k*, and *v_mk_* its standard error associated. The random-effect model incorporates the heterogeneity of the marker effects across studies. Denoting *β_m_* ≔ (*β_m_*_1_*,…, β_mK_*) the vector of the estimated effect of marker *m*, one has

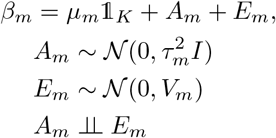

where 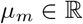 quantifies the deviation to *H*_0_ of marker *m* and is common to all studies, *τ_m_* is between-study variance associated to the random marker effect *A_m_*, *E_m_* the vector of error terms, *V_m_* the diagonal matrix of 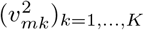 and *I* the identity matrix.

##### Inference

The maximum likelihood estimators of *μ_m_* and 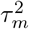 are obtained by solving the following equations iteratively [27]:

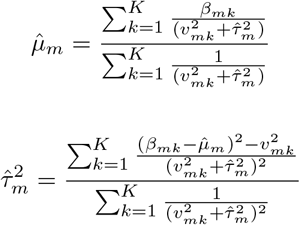

The test of association of the marker corresponds to:

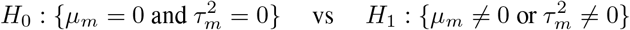

and can be performed using a likelihood ratio test. Noting *l*_0_ and *l*_1_ the likelihood of *β_m_* under *H*_0_ and *H*_1_, respectively, the test statistic is:

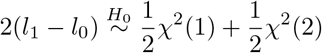

### 4.2 Meta-analysis for G×E analysis in plant genetics

This section introduces the newly developed MA approaches, in the context of plant genetics. The study of G×E interactions in plant genetics requires the evaluation of a same panel - or highly related panels including several common genotypes - in different locations and under different environmental conditions. The use of related panels in MET designs make the independence assumption for the z-scores (see Section 4.1) unrealistic. Depending on the experiment, environments may correspond to controlled stress conditions (*e.g*. nitrogen, water or competition stress), or to different fields and/or years where the environmental conditions are contrasted but not fully controlled by the experimenter. In this section, we show how the fixed and random MA procedures can be adapted to cope with the controlled and uncontrolled environment cases respectively. Assuming that a GWAS analysis has been performed in each environment, the goal of the MA procedure is to summarize the environment-by-environment GWAS results, while i) efficiently controlling the false positive detection rate, and ii) accounting for the heterogeneity of the QTL effects across environments.

#### G×E fixed effect procedure

When the environmental conditions are controlled, environments can be *a priori* classified into several groups. One can then assume the marker effect to be stable within each group, but different from one group to another.

#### Model

We use the same notations as in section 4.1, with *K* now corresponding to the number of environments. We consider that the environments are classified into *J* distinct groups. Since a same panel of varieties - or overlapping panels - are used in all environments, the *Z*-scores cannot be assumed to be mutually independent anymore. The model is updated as follows:

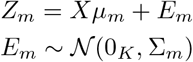

where the incidence matrix 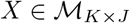 is such that:

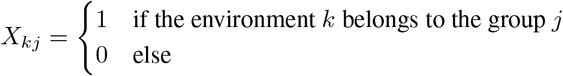

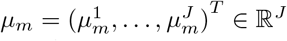 is the vector containing the group specific marker effects, and Σ_m_ is the inter-environment correlation matrix.

Note that if allelic effects are assumed to be stable across all environments, these environments can be gathered into a single group. In such a case the null hypothesis will be *H*_0_: *μ_m_* = 0 as in the classical FE procedure described in Section 4.1.

#### Inference

The parameters to be estimated are the inter-environment correlation matrix Σ_*m*_ and the marker effects within groups 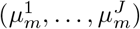. We assume the inter-environment correlation matrix to be common to all markers, i.e. Σ_*m*_ = Σ. Considering only the *M*_0_ markers under *H*_0_, *i.e*. the markers having no effect in any environment, the correlation between the z-scores of two environments *k* and *k′* can be estimated as follows:

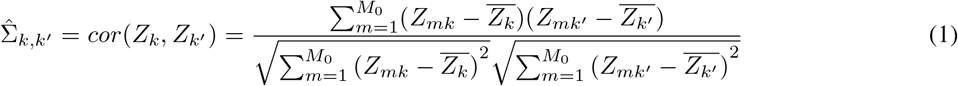

As the list of markers under *H*_0_ is unknown, a filtering step for markers with a high probability of being under *H*_1_ is needed. Different filtering approaches exist, and we present two of them. The first one consists in considering only the markers *m* whose p-values *p_mk_* are higher than a certain threshold fixed by the user, in each environment *k* ∈ 1*,…, K* [47].

The second filtering approach consists, for each environment *k*, in estimating the distribution of the random variables Φ^*−*1^(*p_mk_*), *m* ∈ 1*,…, M*, where Φ^*−*1^ is the inverse distribution function of the normal distribution 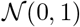. By definition, the distribution to be estimated is a mixture between a 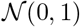 distribution corresponding to the markers under *H*_0_ and a second unknown distribution corresponding to those under *H*_1_. This mixture distribution can be inferred by a kernel method. Then, the filtering consists in considering only the markers *m* whose a posteriori probabilities of being under *H*_1_ are lower than a certain threshold [40]. For the applications detailed in the Section 2, the filtering method was performed with a threshold fixed at 0.8.

Given an estimate of Σ, the within group marker effects 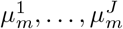 can be inferred by the empirical means of the z-scores of each group:

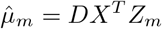

where 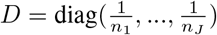 and *n_j_* is the number of experiments in group *j*.

#### Global test

Association of marker *m* can be tested as follows:

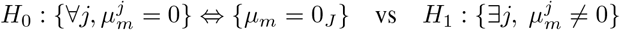

Alternatively, one may test whether the marker *m* has different effects across groups of environments:

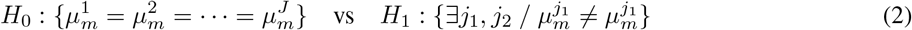

More generally, one may test *H*_0_: {*Cμ_m_* = 0} for any contrast matrix 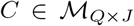, where *Q* ∈ {1, *…, J*} is the number of linear constraints on the marker effects to be tested. Under the null, 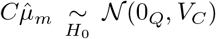. where V_C_ = *CDX^T^ΣX DC^T^* Therefore:

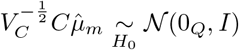

which leads to the following test statistic:

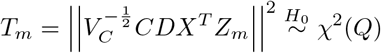

The p-value is given by:

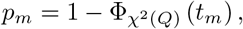

with *t_m_* the observed value of *T_m_*, and 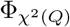 the cumulative distribution function of the *χ*^2^(*Q*) distribution.

#### G×E random effect procedure

In the case of uncontrolled environmental conditions, the heterogeneity of the QTL effects across environments can be accounted for through a random marker effect.

#### Model

The model is updated as follows:

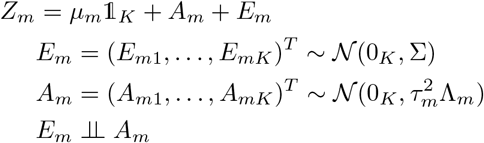

where 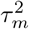 and Λ_*m*_ are the variance and the correlation matrix associated to the random marker effect, respectively.

Compared to the method presented in section 4.1, note that our random effect model is based on the z-scores rather than on the marker effects. As for matrix Σ, we assume the random effect correlation matrix to be common to all markers, *i.e*. Λ_*m*_ = Λ. Furthermore, since Σ quantifies the similarity between environments, it provides some *a priori* knowledge about the similarities of allelic effects at the scale of a marker. Consequently it will be assumed that Λ = Σ.

#### Inference

The correlation matrix Σ is inferred using estimator 1. The effect of the marker 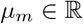 and the between environment variance 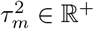 are inferred by maximum likelihood inference, yielding

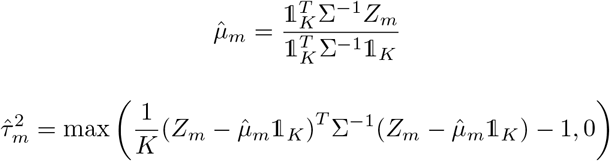

#### Global test

The test for the marker association corresponds to:

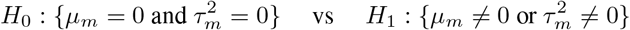

and can be performed using a likelihood ratio test. Noting *l*_0_ and *l*_1_ the likelihood of *Z_m_* under *H*_0_ and *H*_1_, respectively, the test statistic is:

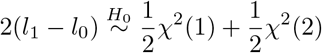

#### Meta-regression test

In MET studies, trials can be characterized through some quantitative environmental covariates (*e.g*. temperature or evapotranspiration). One can then aim at identifying markers whose effects are correlated to a given environmental covariate. Denote the covariate by *X*, and assume *X* to be centered. For each marker *m*, one can test:

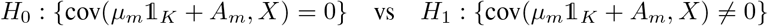

The covariance of interest can be inferred by its empirical counterpart, leading to the following test statistic:

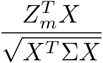

that follows a 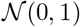 distribution under the null hypothesis.

The p-value is given by:

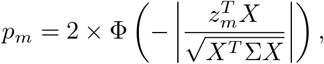

where *z_m_* is the observed value *Z_m_*.

#### Multiple test control

In order to control Type I errors, we apply the local score approach developed by [21]. The local score approach detects significant regions in a genome sequence by accumulating single marker p-values while controlling the FDR. This approach depends on the choice of the threshold *ξ* (in log_10_ scale) below which small p-values are accumulated. The significant genomic regions are computed chromosome by chromosome, and a significance threshold for the FDR control is associated with each chromosome (see [21] for details). For the applications detailed in the Section 2, we set the threshold *ξ* = 3 and fix the nominal FDR level at 0.05.

### 4.3 Availability of data

The raw Arabidopsis data of Frachon et al. (2017) [23] used to perform the individual GWAS is available at the open access local repository https://lipm-browsers.toulouse.inra.fr/pub/Frachon2017-NEE/.

The raw DROPS-Amaizing data of Millet et al. 2019 [42] used to perform the individual GWAS is available at https://doi.org/10.15454/IASSTN.

The raw EU-NAM Flint data of Garin et al. 2020 [24] used to perform the individual GWAS is available at the GitHub repository https://github.com/vincentgarin/mppG×E_data/tree/master/data.

The raw Wheat data of Rincent et al. 2020 [48] used to perform the individual GWAS is available at https://doi.org/10.15454/TKMGCQ.

## 5 Supplementary Material

### A Wheat dataset analysis

We consider the Wheat dataset of [49], where GWAS analyses were performed on a panel of 210 wheat lines phenotyped for grain yield in 16 environments (combinations of location x year x treatment). Lines were genotyped at 108,410 SNPs (after quality control) and phenotyped for heading date and grain yield. In [49] the 16 environments were clustered into 4 groups corresponding to contrasted relationships between heading date and grain yield (low, medium or high correlation between heading date and grain yield, the last group corresponding to a quadratic relationship between heading date and grain yield). When present, the correlation between heading date and grain yield could be positive or negative, depending on the environments. Both the FE and RE were run on the initial per environment GWAS summary statistics.

The RE procedure identified 15 QTLs (Figure.S1.A) each involving one marker except for one QTL located on chromosome 4D involving three markers. Three of these regions were located at less than 1.6 Mb to known flowering genes or heading date QTLs detected on the same panel in [55]. Five of the detected regions colocalized (less than 2Mb) with yield components QTLs detected in [55]. For all these collocalizations, the tests were much more significant with our approach than those from [55] based on a standard GWAS model.

The FE procedure was also applied to detect markers with stable effect across environments. In total 11 QTLs were identified, highlighting some very significant association peaks, especially on chromosome 1D (involving 16 markers), chromosome 6B (involving 13 markers) and chromosome 7B (involving 19 markers) (Figure.S1.B). As expected because of the choice of the 16 environments, these stable regions did not collocalize with major flowering genes nor with heading date QTLs from [55]. Interestingly, two of the detected regions collocalized with yield components QTLs detected in [55], and again the significance was much higher with our approach.

A meta-regression test was performed to detect markers with effects correlated to the correlation between heading date and grain yield (cor_HD_GY). The procedure identified 6 QTLs with two main association peaks on chromosomes 2B and 6A, these regions being illustrated in Fig.S1.C and Fig.S1.D. Two of these six regions were close to a known flowering gene (Ppd-B1 on 2B).

### B Table S1 (Arabidopsis dataset)

### C Night temperature meta-regression tests (Maize dataset)

Regarding night temperature during the grain filling period, hot conditions may affect ovary development and grain growth, carbon translocation or photosynthesis. Interestingly rapid senescence was observed in heat scenarios as well as smaller individual grain sizes and reduced number of grains per ear in the two environments with extreme conditions (Cra). The QTL2_234 found on chromosome 2 was also identified in platform experiments ([45], Granato 2022 in prep) and corresponds to a QTL of plant growth rate in well-watered conditions (pos234) and radiation-interception efficiency (pos233.9).

### D Allelic effects of QTL1_117.6 (EU-NAM Flint dataset)

**Figure S1:**
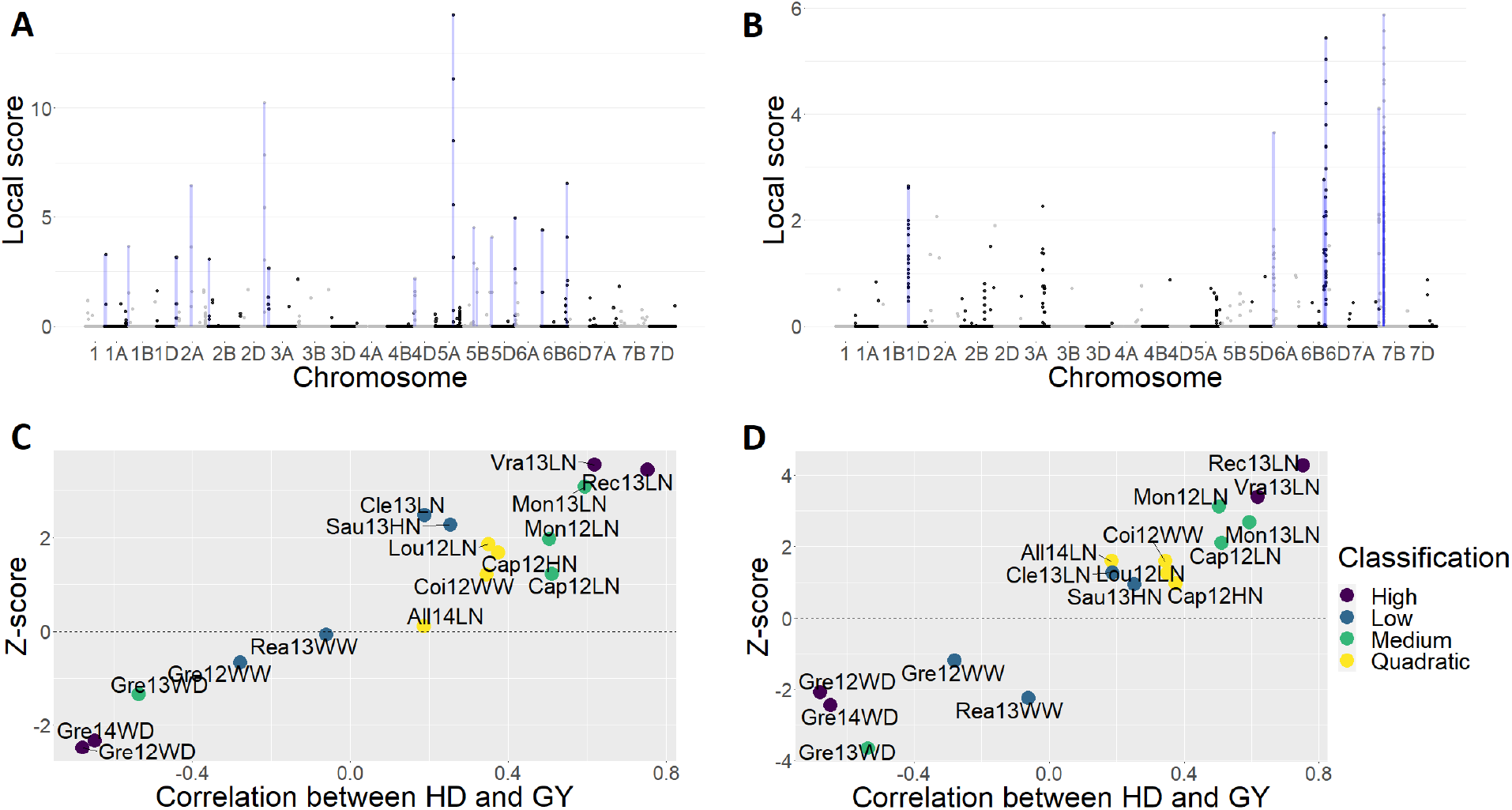
Results of the MA applied to the Wheat dataset. (A) Local score along the chromosomes from the RE procedure. The boxes represent the significant zones identified. (B) Local score along the chromosomes from the FE procedure. The boxes represent the significant zones identified. (C) Z-scores as a function of the correlation between the heading date and the grain yield, for the top significant marker detected with the meta-regression procedure (Wheat dataset, marker cfn2941229 located on chromosome 6A). (D) Z-scores as a function of the correlation between the heading date and the grain yield, for the second top significant marker detected with the meta-regression procedure (Wheat dataset, marker cfn1693678 located on chromosome 2B). Colors correspond to the environment classification according to the relationship between heading date and grain yield of [49].

**Figure S2:**
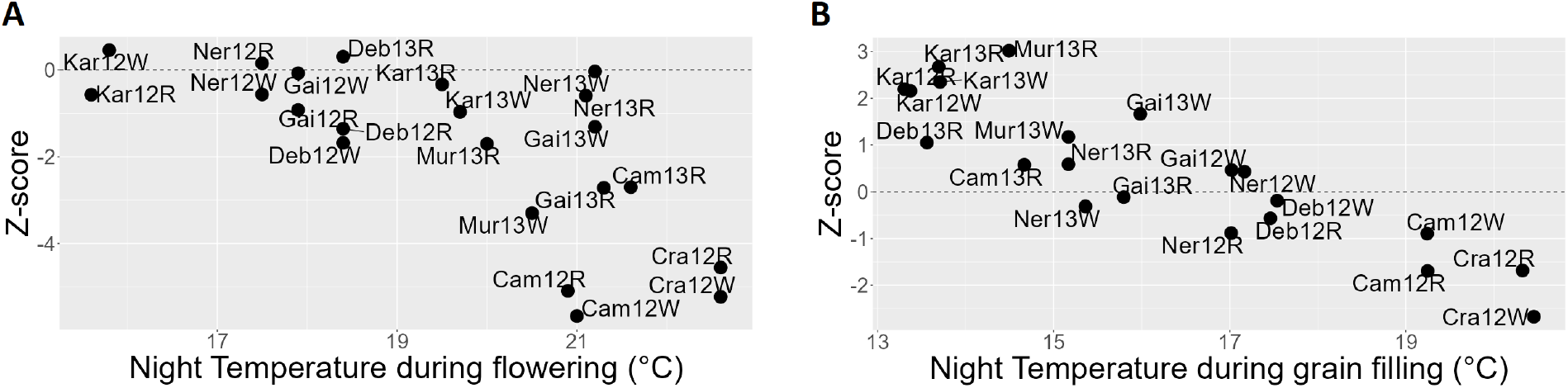
(A) Z-scores as a function of the mean night temperature, for the top significant marker detected with the meta-regression procedure (Maize dataset, marker AX-91369217 located on chromosome 6 pos 21). (B) Z-scores as a function of the mean night temperature during the grain filling period for the top significant marker detected with the meta-regression procedure (Maize dataset, marker AX-91123283 located on chromosome 9 pos 28).

**Figure S3:**
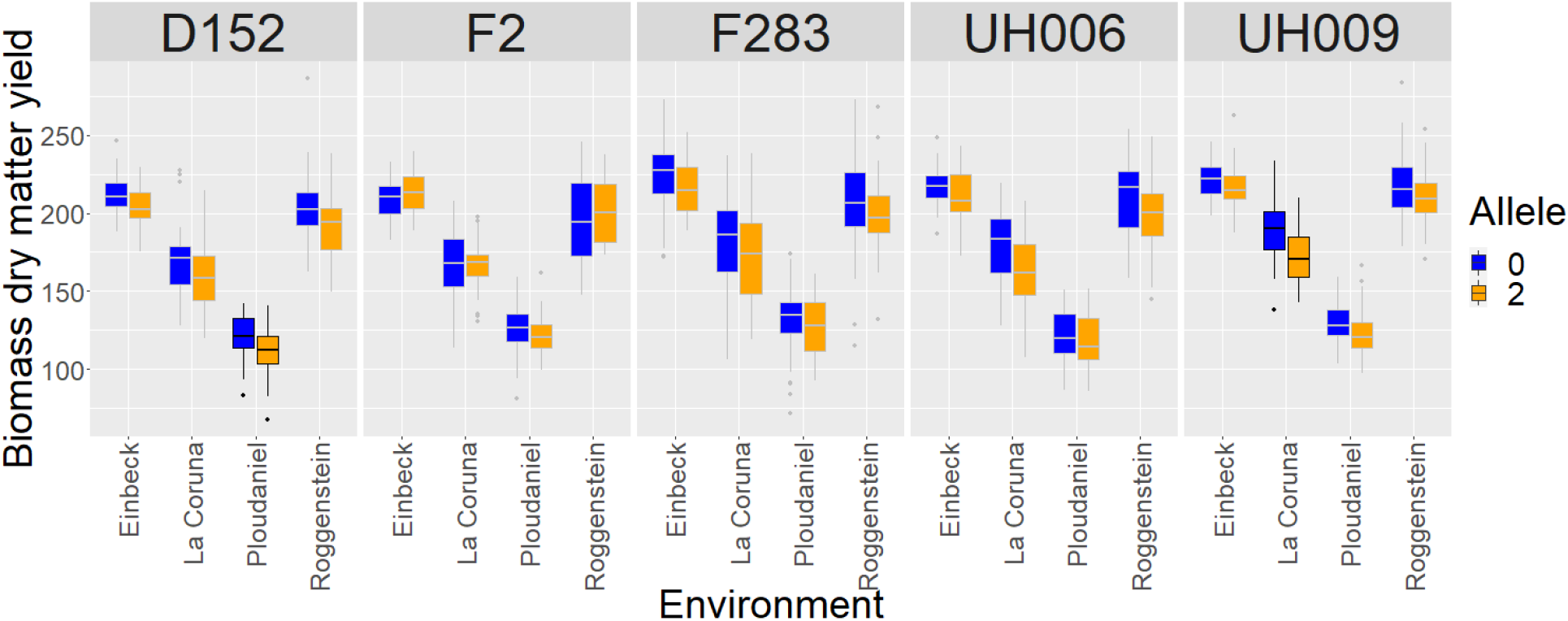
Allelic effects of QTL1_117.6 (EU-NAM Flint, marker PZE.101144585) through locations and sub-populations. A combination of location x sub-population is highlighted if individual GWAS p-value were below 0.01.

**Table S1:** Number of QTLs identified by the standard FE procedure that were found significant in individual analyses. (Arabidopsis dataset analysis)

| Nb. QTLs | Nb. Env. |
| --- | --- |
| 21 | 1 |
| 19 | 2 |
| 11 | 3 |
| 0 | 4 |

